# Strategies for addressing pseudoreplication in multi-patient scRNA-seq data

**DOI:** 10.1101/2024.06.15.599144

**Authors:** Milan Malfait, Jeroen Gilis, Koen Van den Berge, Alemu Takele Assefa, Bie Verbist, Lieven Clement

## Abstract

The rapidly evolving field of single-cell transcriptomics has provided a powerful means for understanding cellular heterogeneity. Large-scale studies with multiple biological samples hold promise for discovering differentially expressed biomarkers with a higher level of confidence through a better characterization of the target population. However, the hierarchical nature of these experiments introduces a significant challenge for downstream statistical analysis. Indeed, despite the availability of numerous differential expression methods, only a select few can accurately address the within-patient correlation of single-cell expression profiles. Furthermore, due to the high computational costs associated with some of these methods, their practical use is limited.

In this manuscript, we undertake a comprehensive assessment of different strategies to address the hierarchical correlation structure in multi-sample scRNA-seq data. We employ synthetic data generated from a simulator that retains the original correlation structure of multi-patient data while making minimal assumptions, providing a robust platform for benchmarking method performance. Our analyses indicate that neglecting within-patient correlation jeopardizes type I error control. We show that, in line with some previous reports and in contrast with others, Poisson Generalized Estimation Equations provide a useful and flexible framework for addressing these issues. We also show that pseudobulk approaches outperform single-cell level methods across the board. In this work, we resolve the conflicting results regarding the utility of GEEs and their performance relative to pseudobulk approaches. As such, we provide valuable guidelines for researchers navigating the complex landscape of gene expression modeling, and offer insights on choosing the most appropriate methods based on the specific structure and design of their datasets.

## Introduction

The cost of single-cell RNA-sequencing (scRNA-seq) technologies has been continuously decreasing and experiments that profile millions of cells are becoming more and more common. In particular, the number of experiments with multiple biological repeats, such as patients from different disease stages, multiple cell lines and experiments with multiple stimuli, is on the rise [1]. Such experiments allow for a better characterization of the target population and are hence pivotal for the discovery of reliable biological markers, i.e., genes that are differentially expressed between groups of subjects [2].

However, such experiments pose new challenges for the statistical analysis. When multiple cells from the same subject are profiled, the cell-level expression profiles can no longer be considered independent. Indeed, expression profiles of cells from the same subject are expected to be more similar than those from cells from different subjects. This within-subject cell-cell correlation, also referred to as pseudoreplication, must be accounted for in the downstream statistical inference [3–5]. However, most current differential expression (DE) methods for single-cell data ignore the hierarchical correlation structure in designs with multiple technical and biological replicates, which eventually results in an accumulation of false positive marker genes.

We illustrate the issues of applying traditional DE methods using a null simulation data set where subjects from the same treatment group are randomly assigned to one of two mock groups. In this setting, genes are not differentially expressed by design, and the DE inference should result in uniformly distributed p-values. However, methods that analyze the data at the single-cell level display an inflation of small p-values, leading to false positive marker gene identifications (Fig 1). For more information on the null simulation datasets, we refer to Section Mock simulation.

**Fig 1.**
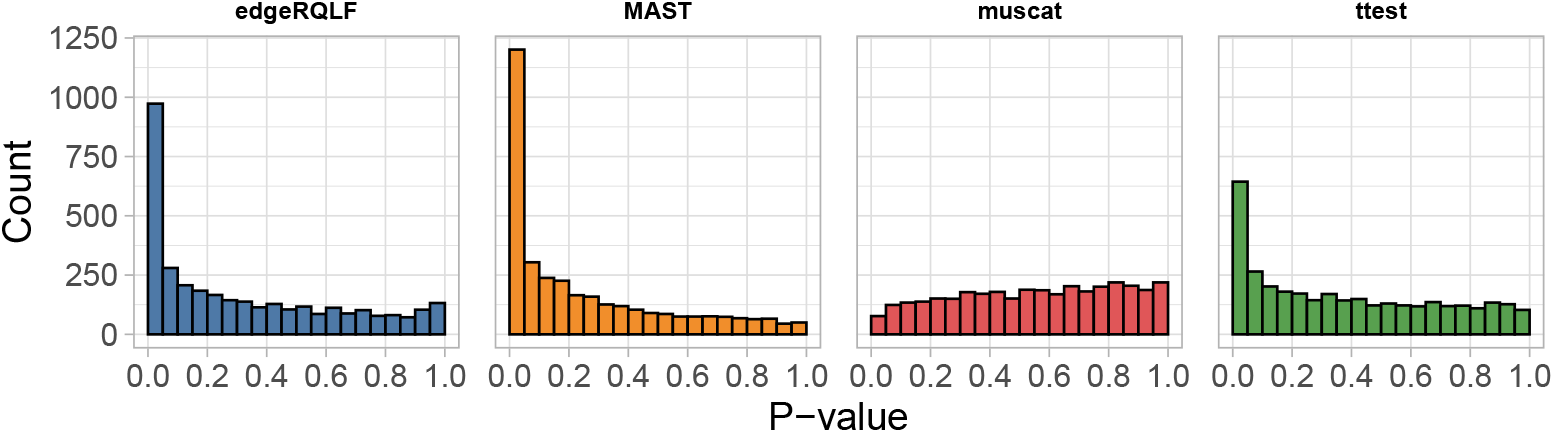
Null simulation results for existing DE methods. DE analysis on the Ledergor null simulation data, where 10 subjects were randomly assigned to one of two mock groups (5 vs. 5 comparison). No systematic differences are expected between the two groups and p-values should be uniformly distributed.

Only *muscat* [4] does not suffer from an increased type I error rate (false positives) in our null simulations. Instead of modeling the data on the single-cell level, *muscat* aggregates the data by taking the sum of the gene counts of all cells from a given subject within a cell type. The resulting observations, i.e., cell type and patient combinations, are referred to as pseudobulk samples. Next, traditional bulk RNA-seq DE methods are applied to the pseudobulk data [4]. As the pseudobulk data only consists of gene abundance measurement per cell type per subject, there is no issue with pseudoreplication when testing for DGE within cell types between sample groups, provided that the different cell types are modeled separately.

However, pseudobulk-level analyses may have several drawbacks. First, by aggregating the data, DE is no longer being evaluated at the single-cell level, thus losing biological resolution. Secondly, it is possible that there are different levels of dispersion between cell types. To allow for a cell-type specific dispersion, and to avoid hierarchical correlation between cell types of the same sample, *muscat* performs a DE analysis for each cell type separately. Hence, the current implementation of *muscat* cannot be used to test for DE between cell types, nor for assessing cell type and treatment interactions, i.e., to test if the treatment effects are changing between cell types.

The statistical power of pseudobulk methods has been subject of active debate in recent literature. On the one hand, Zimmerman *et al*. (2021) [6] claim that pseudobulk methods are underpowered and instead recommend using mixed models with a random effect for subject. Murphy and Skene (2022) [7] revisited the benchmarks of Zimmerman *et al*. (2021) [6] by assessing the type I and type II error rates jointly. They show that pseudobulk methods are in fact the best performers in their updated simulations. This is in line with an extensive benchmark that used real bulk data sets as ground truth [5]. This benchmark presents pseudobulk methods as the best performers and explains the discrepancy in the results between Crowell *et al*. (2020) [4] and Zimmerman *et al*. (2021) [6] by their use of distinct simulation frameworks. The benchmark by Zimmerman *et al*. (2021) [6] also included one additional single-cell level DE modeling strategy. Generalized Estimating Equations (GEEs) are marginal models that address the pseudoreplication problem by explicitly modeling the within-subject correlation in the statistical inference. GEEs can be fitted relatively efficiently and can accommodate count distributions such as the Poisson or negative binomial (NB) distribution, which are commonly used for modeling scRNA-seq data. Zimmerman *et al*. (2021) [6] suggested that GEEs lack performance. However, they used a Gaussian GEE on log-transformed data, which fails to capture the discrete nature of count data. This is expected to be problematic for scRNA-seq, which produces very sparse gene abundance measurements. Due to this sparsity, normalization with logarithmic transformation strongly distorts the characteristics of the original data [8].

In this manuscript, we aim to adequately benchmark existing single-cell level methods, pseudobulk methods and GEEs for multi-subject DE analyses on scRNA-seq data. To allow for a fair comparison, we included GEEs with a Poisson or Negative Binomial variance function and propose a simulation framework with minimal assumptions.

## Materials and methods

### DE methods from the existing literature

In this section, we briefly describe four DE methods from the existing literature that were included in our DE method benchmark. These four strategies are edgeRQLF [9, 10], *muscat* [4], MAST [11] and the Welch t-test [12].

### edgeRQLF

Originally developed for bulk RNA-seq DE analysis, *edgeR* [9, 10] *is currently one of the most popular DE methods for scRNA-seq data. We will use the default edgeR* workflow, which we refer to as *edgeRQLF. edgeRQLF* models *Y*_*gij*_, i.e., the expression count of gene *g* in cell *j* within subject *i*, using the following quasi-negative binomial Generalized Linear Model (GLM):

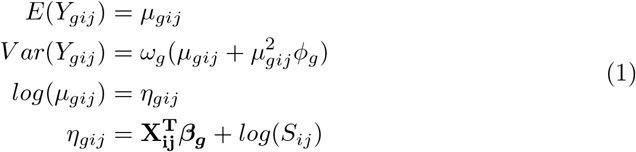

where *µ*_*gij*_ is the unobserved average expression of gene *g* in cell *j* of subject *i, φ*_*g*_ is the gene-level dispersion parameter of the negative binomial model and *ω*_*g*_ the quasi-likelihood dispersion parameter. ***β***_***g***_ is a *p ×* 1 column vector of regression parameters modeling the association between the average gene counts and covariates 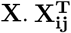 is a row in the *N ×*p matrix of covariates **X**, which corresponds with the covariate pattern of cell *j* from subject *i*. In addition, a normalization factor or size factor *S*_*ij*_ is included in the linear predictor, which is used to correct for technical differences between cells, i.e., differences in the library size (total number of sequenced reads per cell) and RNA composition. For this, *edgeR* by default uses trimmed-mean of M-values (TMM) normalization factors [13]. Inference on the regression parameters ***β***_***g***_ is achieved using an F-test.

Note that the GLM framework used by *edgeRQLF* cannot readily address the hierarchical correlation structure observed in multi-sample scRNA-seq datasets. When the hypothesis involves an effect that can be estimated within each sample, e.g., studying DGE between cell types, *edgeRQLF* can adequately estimate this effect by including a fixed effect for sample in the matrix of covariates **X**_**ij**_ (blocking). However, when the hypothesis requires effects that cannot be estimated within each block, e.g., DGE for each cell type between sample groups, then the correlation structure in the data cannot be addressed using sample-level blocking. In such settings, *edgeRQLF* does not adequately address the hierarchical correlation structure in the data, which may eventually result in an accumulation of false positive marker genes.

### muscat

*muscat* [4] builds upon the modeling framework of *edgeRQLF*. The main difference with *edgeRQLF* is that *muscat* first aggregates the single-cell level RNA-seq data to the “pseudobulk” level. In the default implementation, *muscat* generates pseudobulk samples by summing the gene expression counts for cells within each “cell type - subject” combination. Subsequently, the aggregated counts are modeled per cell type in the same way as described for *edgeRQLF* (Eq 1).

One important consequence of pseudobulk aggregation is that the number of repeated measurements within a subject is greatly reduced. In the single-cell level data, the number of repeated measurements is the total number of cells sequenced for subject *i, n*_*i*_, which can be thousands of cells. By contrast, in pseudobulk data, the number of measurements per subject equals the number of unique cell types identified in that subject, a number that typically is well below 50.

As discussed above for *edgeRQLF* (Section edgeRQLF), *muscat* also cannot readily accommodate the hierarchical correlation structure observed in multi-sample scRNA-seq data. As such, the same concerns about pseudoreplication apply here. However, note that as the pseudobulk data only consists of gene abundance measurement per cell type per subject, the number of repeated measurements for each sample is greatly reduced. Furthermore, the default implementation of *muscat*, which tests for DGE within cell types between sample groups, will analyze each cell type separately. As such, these analyses no longer have multiple measurements per sample.

### MAST

MAST [11] is another popular DE method for scRNA-seq data analysis. MAST first normalizes the scRNA-seq data to transcript-per-million (TPM) abundances, i.e., by (1) dividing the gene counts by the length of the gene in kilobases, (2) dividing this by a cell specific scaling factor that is computed by taking the sum across the abundances obtained in step (1), and (3) multiplying by one million. Next, MAST models the normalized expression 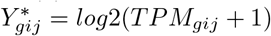 of gene *g* in cell *j* of subject *i* with a two-part generalized regression model, also referred to as a hurdle model. The first component consists of a logistic regression model on the binarized expression matrix, which can make inference on differential detection between conditions. The second component models the gene expression of cells for which the gene has positive counts, assuming a truncated Gaussian model for 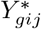. This model component can be used to infer DE. Furthermore, Finak *et al*. (2015) [11] use the cellular detection rate, i.e., the fraction of genes detected in each cell, as an additional covariate in both model components to normalize for between-cell differences due to technical variability, e.g. mRNA quality, pre-amplification efficiency of the scRNA-seq assay, and nuisance biological variability such as differences in cell volume. A hurdle model implies estimation orthogonality. As such, the detection and expression components can be modeled and tested separately, i.e., using a likelihood ratio test or a Wald test. Upon summation, the test statistics remain asymptotically 𝒳^2^ under the null hypothesis.

### Welch t-test

For comparison, we also apply a classical Welch t-test [12]. More specifically, we use the *pairwiseTTests* function from the *scran* package to model 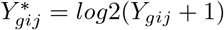 expression counts that are additionally normalized for differences in the sequencing depth. This function allows for blocking on uninteresting factors of variation, e.g., the different sequencing batches of the Lupus dataset (Section Mock simulation), by performing Welch t-tests between biological groups within each block. The p-values for each gene across the different blocks are combined using Stouffer’s weighted Z-score method [14]. The reported log-fold change for each gene is also a weighted average of log-fold changes across blocks.

### Generalized Estimation Equations for multi-patient scRNA-seq DE testing

In this section, we describe Generalized Estimation Equations (GEEs) in the context of DE testing for multi-patient scRNA-seq data. First, we specify our vanilla GEE implementation. Subsequently, we discuss different variations to this vanilla implementation, including deviations from the mean model component specification, deviations from the variance specification, a small sample size correction strategy and a deviation in the inference method. Finally, we provide Table 1, which serves as a summary of all DE methods that were benchmarked in this manuscript, including all GEE variations and four methods from the existing literature. For a more detailed introduction to GEEs and related concepts, including the “sandwich” estimator, we refer to the technical note in Supporting Information.

**Table 1.** Overview of the different DE methods included discussed and evaluated in this manuscript. Column 1: Name. Identifier of the DE method. These identifiers will be used to refer to a method in the text and in each figure legend. **Column 2: Description**. Link to the section where we provide a more detailed description of the DE method. **Column 3: Pseudobulk**. Indicates if the DE method performs pseudobulk aggregation (yes/no). **Column 4: Normalization**. Indicates the type of data normalization. Either TMM normalization (Section Vanilla GEE for DE testing), logarithmic transformation, or no normalization. **Column 5: Variance function**. Indicates the function used to model the variance in the data. **Column 5: Pseudo-count**. The size of the pseudo-count that was added to the data to stabilize the parameter estimation procedure. **Column 6: Cluster level**. Whether the model specifically models within-subject correlation (*subject/none*). **Column 7: Small corr** Whether a small sample bias correction was applied. **Column 8: Test**. Statistical test used for inference on DGE.

| Name | Description | pseudo-bulk | Normalization | Variance function | Pseudo-count | Cluster level | Small corr | Test |
| --- | --- | --- | --- | --- | --- | --- | --- | --- |
| edgeRQLF | link1 | No | TMM | Negative-Binomial | 0.125 | None | No | F-test |
| *muscat* | link2 | Yes | TMM | Negative-Binomial | 0 | None | No | F-test |
| MAST | link3 | No | Log | Gaussian | 0 | None | No | Z-test |
| t-test | link4 | No | None | Gaussian | 0 | None | No | T-test |
| sandwich_base | link5 | No | TMM | Poisson | 0 | Subject | No | Z-test |
| sandwich_SS | link5;link8 | No | TMM | Poisson | 0 | Subject | Yes | Z-test |
| sandwich_Ttest | link5;link9 | No | TMM | Poisson | 0 | Subject | No | T-test |
| sandwich_SS_Ttest | link5;link8;link9 | No | TMM | Poisson | 0 | Subject | Yes | T-test |
| sandwich_ps0125_SS_Ttest | link5;link6;link8;link9 | No | TMM | Poisson | 0.125 | Subject | Yes | T-test |
| **sandwich_ps1_SS_Ttest** | link5;link6;link8;link9 | No | TMM | Poisson | 1 | Subject | Yes | T-test |
| sandwich_NB_SS | link5;link7;link8 | No | TMM | Negative-Binomial | 0 | Subject | Yes | Z-test |
| sandwich_NB_SS_Ttest | link5;link7;link8;link9 | No | TMM | Negative-Binomial | 0 | Subject | Yes | T-test |

### Vanilla GEE for DE testing

Generalized Estimation Equations provide a statistical framework for extending GLMs to clustered and longitudinal data [15]. As such, GEEs can acknowledge the hierarchical correlation structure in multi-sample scRNA-seq data by explicitly modeling the within-subject correlation at the single-cell level. As an example, we take an scRNA-seq dataset with multiple subjects and multiple cells per subjects. Without loss of generality, we will consider a single gene *g*, and henceforth drop the subscript *g*. Let:

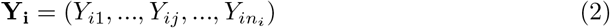

be a *n*_*i*_ *×* 1 column vector of responses (measured gene abundances) for cells *j* = 1, …, *n*_*i*_ of subject *i* = 1, …, *N* and

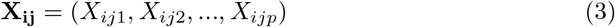

a *p ×* 1 column vector of between-subject and within-subject covariates potentially associated with the response *Y*_*ij*_.

GEEs specify (i) the expectation of the response conditional on the fixed effects 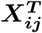, (ii) the conditional variance of *Y*_*ij*_ given 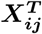 and (iii) the conditional within-subject association among the vector of repeated responses [15, 16], as follows:

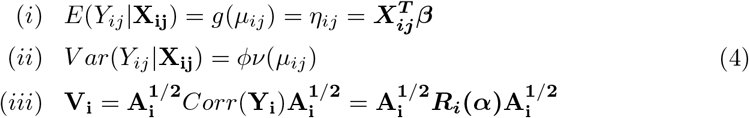

Component (i) relates the expectation of the response with a linear predictor of covariates through a known link function *g*(.). Note that this specification coincides with that of the mean component of a canonical GLM. Both GEEs and GLMs estimate population-averaged or marginal effects. Indeed, the estimator of the mean model parameters of the GEE is conditional only on the fixed effects 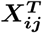, and not conditional on any random effects.

Component (ii) describes the mean-variance relationship, i.e., the relationship between the variance of *Y*_*ij*_ given the covariates **X**_**ij**_ according to a known variance function *ν*_*i*_(.), and an additional scale or dispersion parameter *φ*. Note that in the most general formulation, which is displayed here, a single dispersion parameter *φ* is estimated. In principle, *φ* can be estimated separately for different covariate levels, e.g., *φ* can be cell type specific.

Component (iii) explicitly describes the within-subject correlation conditional on the covariates **X**_**ij**_, as a function of the conditional means *µ*_*ij*_ and an additional set of within-subject association parameters *α*. More specifically, **A**_**i**_ is a diagonal matrix with *V ar*(*Y*_*ij*_|**X**_**ij**_) = *φν*(*µ*_*ij*_) along the diagonal. As such, 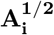 is a diagonal matrix of the standard deviations. *Corr*(**Y**_**i**_) = ***R*(*α*)** is a matrix describing the correlation among repeated measurements within subject *i* as a function of parameters *α*. In the special case *α* = 0, the repeated measurements within subject *i* are considered independent from one another. Finally, **V**_**i**_ is typically referred to as the “working” covariance matrix, which is allowed to be misspecified, as shown the technical note in Supporting Information.

It is important to note that a conventional GLM with a sandwich estimator is a special case of a GEE. Indeed, for GEEs that use an independent correlation structure (see technical note in Supporting Information), the estimation of the mean model parameters reduces to a Newton-Raphson algorithm that is analogous to that of a traditional GLM. Subsequently using sandwich estimator to obtain the variance makes the inference robust towards the misspecification of the working correlation. This analogy can be readily derived from Eq 4. Indeed, when assuming an independence working correlation structure and a Poisson variance structure, component (ii) of Eq 4 reduces to *V ar*(*Y*_*gij*_|**X**_**ij**_) = *φν*(*µ*_*gij*_) = *φµ*_*gij*_, and component (iii) of Eq 4 reduces to **V**_**i**_ = **A**_**i**_. In the remainder of this manuscript, we henceforth refer to GEEs with an independent correlation structure as “sandwich”.

As such, we can specify a Poisson GLM for gene *g* ∈ {1, …, *G*} as

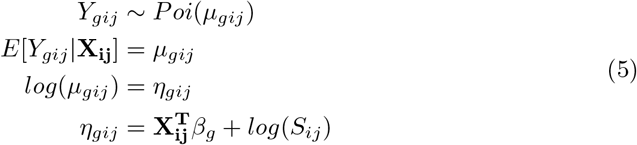

to model the expression *Y*_*gij*_ of gene *g* in cell *j* = 1, …, *n*_*i*_ of subject *i* = 1, …, *N*. ***β***_***g***_ is a *p ×* 1 column vector of regression parameters modeling the association between the average gene counts and the covariates **X**. Finally, 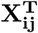 is a row in the *N ×*p design matrix **X** that corresponds with the covariate pattern of subject *i* cell *j*, with *p* the number of parameters of the mean model, i.e., the length of vector ***β***_***g***_. In addition, a normalization factor or size factor *S*_*ij*_ is included in the linear predictor, which is used to correct for technical differences between cells, i.e., differences in the library size (total number of sequenced reads per cell) and RNA composition. For this, we use the trimmed-mean of M-values (TMM) normalization factors from edgeR [13]. We use a sandwich estimator to obtain a robust variance estimate for 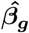 using the *vcovCL* function of the *sandwich* package [17]. The *vcovCL* function allows for acknowledging the within-subject association among the vector of repeated measurements by using an empirical estimator to correct for within-patient correlation. We will refer to the model from Eq 5 as *sandwich_base*.

Next, we infer on (linear combinations of) 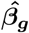 from Eq 5 using a Wald test, with null hypothesis

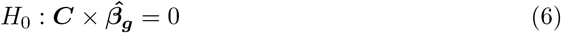

i.e., no shift in the average gene expression between biological groups, and ***C*** the matrix representing the linear contrasts of interest for the DGE test. The Wald test statistics are assumed to asymptotically follow a standard normal distribution under the null hypothesis of no association.

### Adding a pseudo-count

The first deviation from the baseline GEE method described in Section Vanilla GEE for DE testing is to add a pseudo-count to the data. This deviation will affect the specification of the mean model component, i.e.,

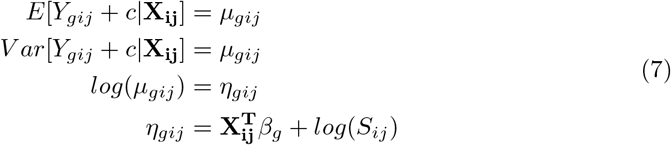

with pseudo-count *c*. The goal of the pseudo-count is to stabilize the parameter estimation procedure when the data are sparse.

There are two models discussed in this manuscript that incorporate a pseudo-count (denoted *ps*), i.e., *sandwich_ps0125_SS_Ttest* and *sandwich_ps1_SS_Ttest*, with a pseudo-count of 0.125 and 1, respectively.

### Deviation in the variance function

The second deviation from the baseline GEE method described in Section Vanilla GEE for DE testing is to assume a Negative Binomial variance function for *Y*_*gij*_. This deviation will affect the specification for the variance of *Y*_*gij*_, i.e.,

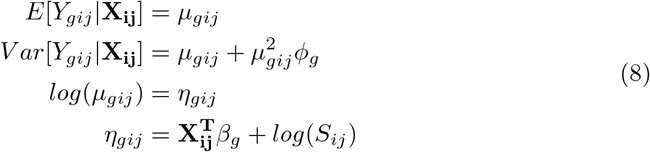

with *φ*_*g*_ the dispersion parameter of the Negative Binomial variance function. As such, this specification allows for modeling a mean-variance trend in the RNA abundances. Subsequently, we again obtain robust sandwich variance estimates for 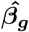 using the *vcovCL* function of the *sandwich* package [17]..

Two models discussed in this manuscript adopt the Negative Binomial variance function (denoted *NB*), i.e., *sandwich_NB_SS* and *sandwich_NB_SS_Ttest*.

### Small-sample bias correction

As pointed out in Section Generalized Estimation Equations for multi-patient scRNA-seq DE testing, the sandwich estimator is robust against a potential misspecification of the “working” covariance matrix **V**_**i**_. However, this robustness property only holds asymptotically, i.e., when (i) the number of subjects *N* is relatively large, (ii) the number of repeated measurements per patient *n*_*i*_ is relatively small and the number of repeated measurements is balanced across subjects. These requirements are typically not met in multi-patient scRNA-seq datasets. Consequently, sandwich-based standard errors can be expected to be biased downward, resulting in overly liberal inference [16]. Since settings where *N* is small and *n*_*i*_ is large are common, a large body of literature exists to improve the finite sample properties of the sandwich estimator [18–22]. **We here adopt the simple small sample size adjustment first proposed** by MacKinnon and White (1985) [19], which multiplies the sandwich variance by 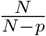, where *N* is the number of clusters and *p* is the number of regression parameters.

Six models discussed in this manuscript adopt this small sample size (denoted *SS*), i.e., *sandwich_SS, sandwich_SS Ttest, sandwich ps0125_SS_Ttest, sandwich_ps1_SS_Ttest, sandwich_NB_SS* and *sandwich_NB_SS_Ttest*.

### Inference with approximate T-test

The baseline GEE method *sandwich_base* achieves inference on 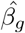 by using a Wald test, where the Wald test statistic is assumed to asymptotically follow a standard normal distribution under the null hypothesis of no association. Here, we evaluate a heuristic workflow where we assume the Wald statistics to follow a T-distribution under the null hypothesis, with degrees of freedom *df* = *N* − *p*_*between*_, with *N* the total number of subjects and *p*_*between*_ the number of between-subject model parameters, i.e., the number of parameters that cannot be estimated within subject.

Five models discussed in this manuscript use a Wald T-test (denoted *Ttest*), i.e., *sandwich_Ttest, sandwich_SS_Ttest, sandwich_ps0125_SS_Ttest, sandwich_ps1_SS_Ttest*, and *sandwich_NB_SS_ Ttest*.

### A summary of all DE models in this study

In Table 1, we provide a schematic overview of the different DE methods included discussed and evaluated in this manuscript.

### Mock simulation

The “Ledergor” dataset is derived from a study on multiple myeloma [23]. The original publication utilized the plate-based scRNA-seq technology *MARS-seq* [24] to sequence 20.568 bone marrow plasma cells (i.e., one single cell type) for 40 individuals along the multiple myeloma progression spectrum (11 control donors, 29 multiple myeloma patients with varying disease severity). We first filtered the data to obtain a biologically homogeneous subset of samples, retaining only the samples from the 11 healthy donors. We adopted the same gene-level filtering strategy as the original publication, removing genes with a mean expression less than 0.001 unique molecular identifiers (UMIs) per cell, and genes with both an above-average expression and a coefficient of variance below a value of 1.2.

Next, the 11 samples were split randomly into two groups of 5 samples each, randomly omitting 1 sample. As we assume these 11 patients to all be members of the same underlying population, i.e., we assume this subset to be biologically homogeneous, this randomization is not expected to be biologically meaningful. Hence, we make the assumption that genes, on average, should not display DE signal between these two random groups, which we thus will refer to as “mock” groups. Note that we repeated this random mock group assignment five times to average out the effect of potential unmeasured covariates, i.e., to reduce the impact of our assumption that these 11 patients are biologically homogeneous.

The “Lupus” dataset is derived from a study on systemic lupus erythematosus (SLE) [25]. The original publication utilized the droplet-based scRNA-seq technology *mux-seq* [26] to profile over 1.2 million peripheral blood mononuclear cells (PBMCs) derived from 162 systemic lupus erythematosus (SLE) cases and 99 healthy controls of Asian or European ancestry. Like with the Ledergor dataset, we first filtered the data to obtain a homogeneous subset retaining only samples from healthy, European women under 50 years old. Additionally, only subjects from three specific sequencing batches were included (batches dmx_count_AH7TNHDMXX_YE_8-30, dmx_count_AHCM2CDMXX_YE_0831 and dmx_count_BH7YT2DMXX_YE_0907), thus retaining 44 patients. To further enhance homogeneity, we analyzed cell types separately, focusing on T4 naive cells, B memory cells, and non-classical myeloid cells. These three cell types were selected because they were observed in all patients, and because they have a high, intermediate and relatively low number of cells per patient, respectively. The median number of cells per patient is 708 cells for the T4 naive cell type, 321 cells for the B memory cell type and 118.5 cells for the non-classical myeloid cells. Gene-level filtering was performed separately for each cell type, only retaining genes that have an expression of at least 10 unique molecular identifiers (UMIs) over all cells, and are expressed in at least 2% of all cells. Next, mock datasets were generated in the same way as described above for the Ledergor dataset, i.e., by randomly splitting the 44 samples into two mock groups. Here, we specifically made sure to have the same number of samples per sequencing batch in both mock groups. Again, the random splitting was repeated five time to reduce the impact of potential unmeasured covariates. Each dataset (cell type) was analyzed separately.

### Simulations with DE signal

We aimed to evaluate the sensitivity and specificity of different DE methods on multi-patient scRNA-seq datasets with ground truth DE signal. However, Crowell *et al*. (2022) [27] argue that there are currently no scRNA-seq simulation workflows available that faithfully capture the within-subject correlation structure and other key characteristics of real data. Therefore, we propose the following workflow.

As a starting point, we use the mock simulation datasets from Section Simulations with DE signal. Next, we randomly swapped gene expression between 5% of the genes in one of the mock treatment groups. This swapping of gene counts introduces a differential gene expression signal of different magnitudes between the mock treatment groups. Treating the swapped genes as truly differential features allows for benchmarking the sensitivity and specificity of the different DE workflows. By starting from a real dataset, we avoid making strong assumptions on the underlying data characteristics (e.g., assuming that the expression of each gene is a realization of a negative binomial distribution). In addition, our simulation workflow in principle should be able to retain the within-subject and between-gene correlation structures from the original data. The implementation of this minimal-assumptions simulator is available from https://github.com/milanmlft/swapper. We used the *countsimQC* package [28] to compare our simulated data to the original Ledergor dataset with regard to several data characteristics (e.g., library size, gene-gene correlation distribution, cell-cell correlation distribution). These *countsimQC* reports indicate that our simulations adequately mimic real data. These reports are available from https://doi.org/10.5281/zenodo.10799944. Simulated datasets were created based on the Ledergor dataset and based on the three cell types (T4 naive cell, B memory cells and non-classical myeloid cells) of the Lupus dataset. Each dataset (cell type) was analyzed separately. Again, each analysis was performed five times using a different randomly selected subset of samples.

### Performance measures

We assess the performance of the different DE methods by visualizing the true positive rate (TPR) against the false discovery rate (FDR), which are computed according to the following definitions:

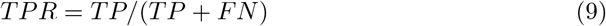

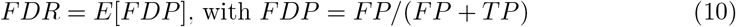

where FN, FP and TP denote the number of false negatives, false positives and true positives, respectively. The TPR is a measure of sensitivity, while FDR is the expected value of the false discovery proportion (FDP), which is a measure of specificity.

## Code and data availability

The source code to reproduce our analyses and figures displayed in this manuscript is available from

https://github.com/statOmics/PseudoreplicationPaper. These repositories contain all information to run our analyses from scratch. For snapshots of these repositories after running all the scripts, containing all input, intermediate data objects and output, we refer to our Zenodo repository https://doi.org/10.5281/zenodo.10821869.

## Results

This results section is organized as follows: we first benchmark the type I error control of the different DE methods using null simulations. Results for the five “main” DE methods, i.e., edgeRQLF, *muscat*, MAST, t-test and sandwich_ps1_SS_Ttest (Table 1), will be displayed in the body of the text, whereas results for all other methods are available as supplementary figures (Supporting Information). Next, we benchmark sensitivity and specificity of the DE methods in simulations with ground truth DE signal, and characterize the effect of sample size on the performance. Finally, we extend our benchmark for sensitivity and specificity to datasets with multiple cell types. All simulations were generated based on two real single-cell transcriptomics datasets, here referred to as the “Ledergor” and “Lupus” datasets, obtained from Ledergor *et al*. (2018) [23] and Perez *et al*. (2022) [25], respectively. For more details on these datasets, we refer the reader to Sections Mock simulation and Simulations with DE signal. Note that the Ledergor dataset was generated using the plate-based scRNA-seq protocol MARS-seq [24], while the Lupus dataset was acquired using the droplet-based scRNA-seq protocol mux-seq [26].

### Mock analysis

We benchmark the type I error control of the different DE methods using null simulations, i.e., simulations in which none of the genes are DE by design. As such, all genes that are flagged as DE between conditions are false positives. The null simulations were generated based on two real single-cell transcriptomics datasets, here referred to as the “Ledergor” and “Lupus” datasets, obtained from Ledergor *et al*. (2018) [23] and Perez *et al*. (2022) [25], respectively. We refer the reader to Section Mock simulation for more details on the design of the null simulation data sets.

In Fig 2, we show the p-value densities of five DE methods on two null simulation data sets. The null simulations in row A of Fig 2 are based on the Ledergor data, which consists of five samples per mock treatment group. The null simulations in row B of Fig 2 are based on the B memory cells from the Lupus data, which consists of twenty-two samples per mock treatment group. For both data sets, the analysis was performed five times, each time with a different randomly selected subset of samples per mock treatment group, as represented by the different line types.

**Fig 2.**
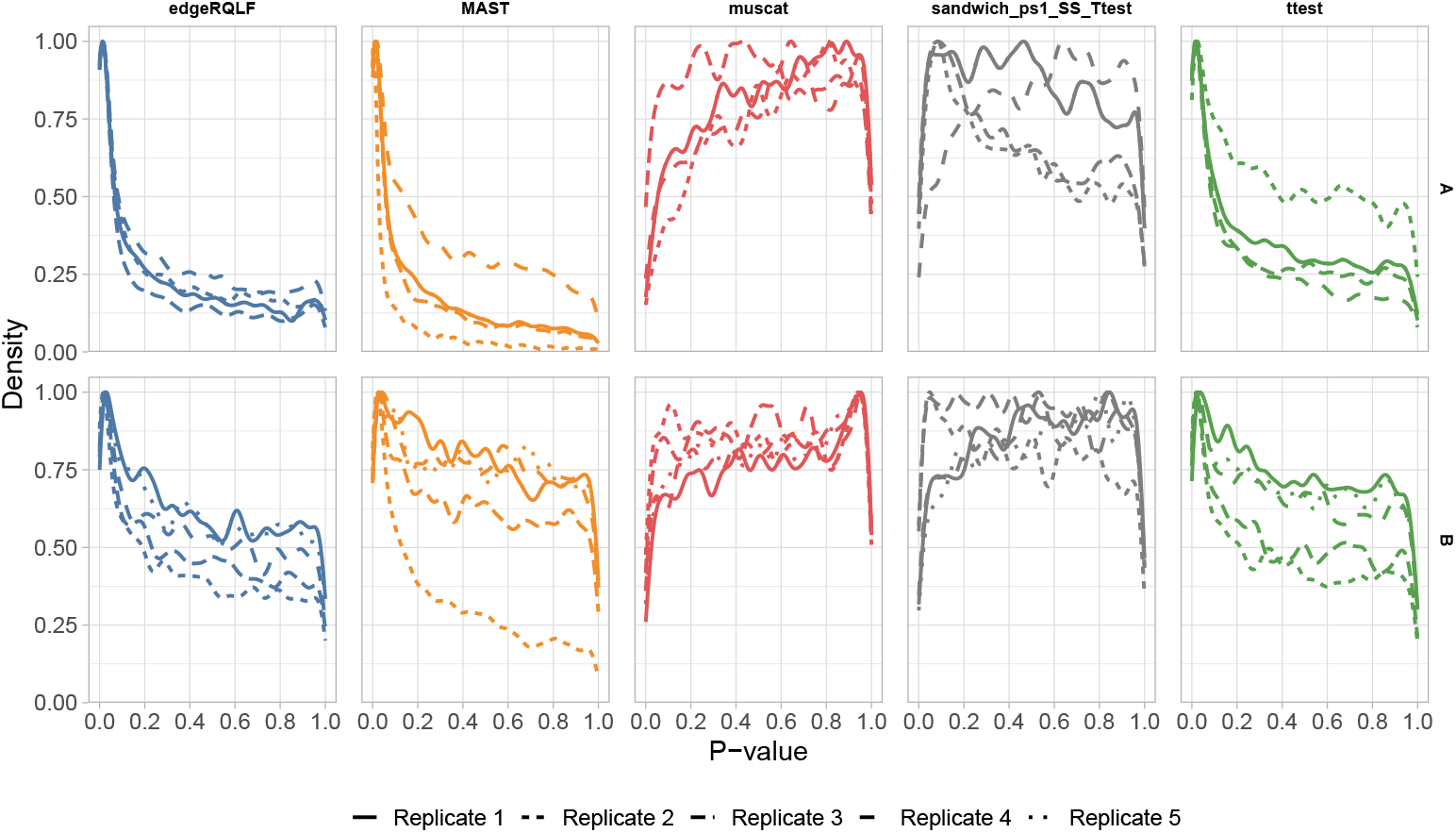
P-value densities of DE methods for multi-subject null simulations. **Row A:** P-value density plots for the different DE methods on null simulations based on the Ledergor data (5 vs. 5 subjects comparison). Compared to 1, this row additionally includes a panel for the sandwich method. The different line types indicate the p-value densities for the five different data replicates. **Row B:** P-value density plots for the different DE methods on null simulations based on the memory B-cells of the Lupus data (22 vs. 22 subjects comparison).

For these null simulations, the p-value densities should follow the uniform distribution when the DE method is controlling the type I error, which would appear in Fig 2 as horizontal lines. The three canonical single-cell methods that ignore pseudoreplication, i.e., edgeRQLF, MAST and ttest, provide a poor control of the type I error across all simulations, indicating an accumulation of false positive results. As already shown in Fig 1, the pseudobulk-level analysis method *muscat* provides better type I error control, because the pseudobulk aggregation removes pseudoreplication altogether when assessing DE within cell type. The sandwich approach, which models the data at the single-cell level but explicitly acknowledges within-sample cell-cell correlation, also controls the type I error reasonably well. Taken together, these results confirm the need of accounting for within-subject correlation when performing DE analyses on multi-subject single-cell data.

P-value density plots on the mock simulation data for all DE methods described in Sections DE methods from the existing literature and Generalized Estimation Equations for multi-patient scRNA-seq DE testing and all datasets (the Ledergor dataset and three cell types from the Lupus dataset) are displayed in Supplementary Fig S1-S4. Across all datasets, methods that model the data at the single-cell level and that do not specifically acknowledge the within-subject cell-to-cell correlation (i.e., edgeRQLF, MAST and ttest) have overly liberal inference. In the evaluation on the Ledergor dataset in Supplementary Fig S1, which has the smallest sample size (5 versus 5 patients), a decrease in type I error control is observed for GEE-based methods that do not perform a small sample size correction in their inference (sandwich base and sandwich Ttest), or that rely on z-test inference rather than t-test inference (sandwich base, sandwich_NB_SS, sandwich_SS). We did not observe a large impact of the pseudo-count used in the GEE-modeling procedure on the type I error control, i.e., sandwich_SS_Ttest, sandwich_ps0125_SS_Ttest and sandwich_ps1_SS_Ttest all adequately controlled the type I error across all datasets.

Note that some methods are excluded from Supplementary Fig S4 due to their long running times. This will be discussed in more detail in Section Scalability assessment.

### Scalability assessment

We evaluate the scalability of all DE methods on the mock simulation datasets based on the Ledergor and the three cell types of the Lupus datasets (see Section Mock simulation). These datasets all have a different number of observations and a different number measurements per observation, i.e., number of gene abundances measured per observation (Ledergor data: 983 observations, 3.411 measurements; Lupus data B memory cell type: 8.724 observations, 4.815 measurements; Lupus data non-classical myeloid cell type: 5.624 observations, 6.081 measurements; Lupus data T4 naive cell type: 36.901 observations, 4.013 measurements). The running time of each method is displayed in Fig 3.

**Fig 3.**
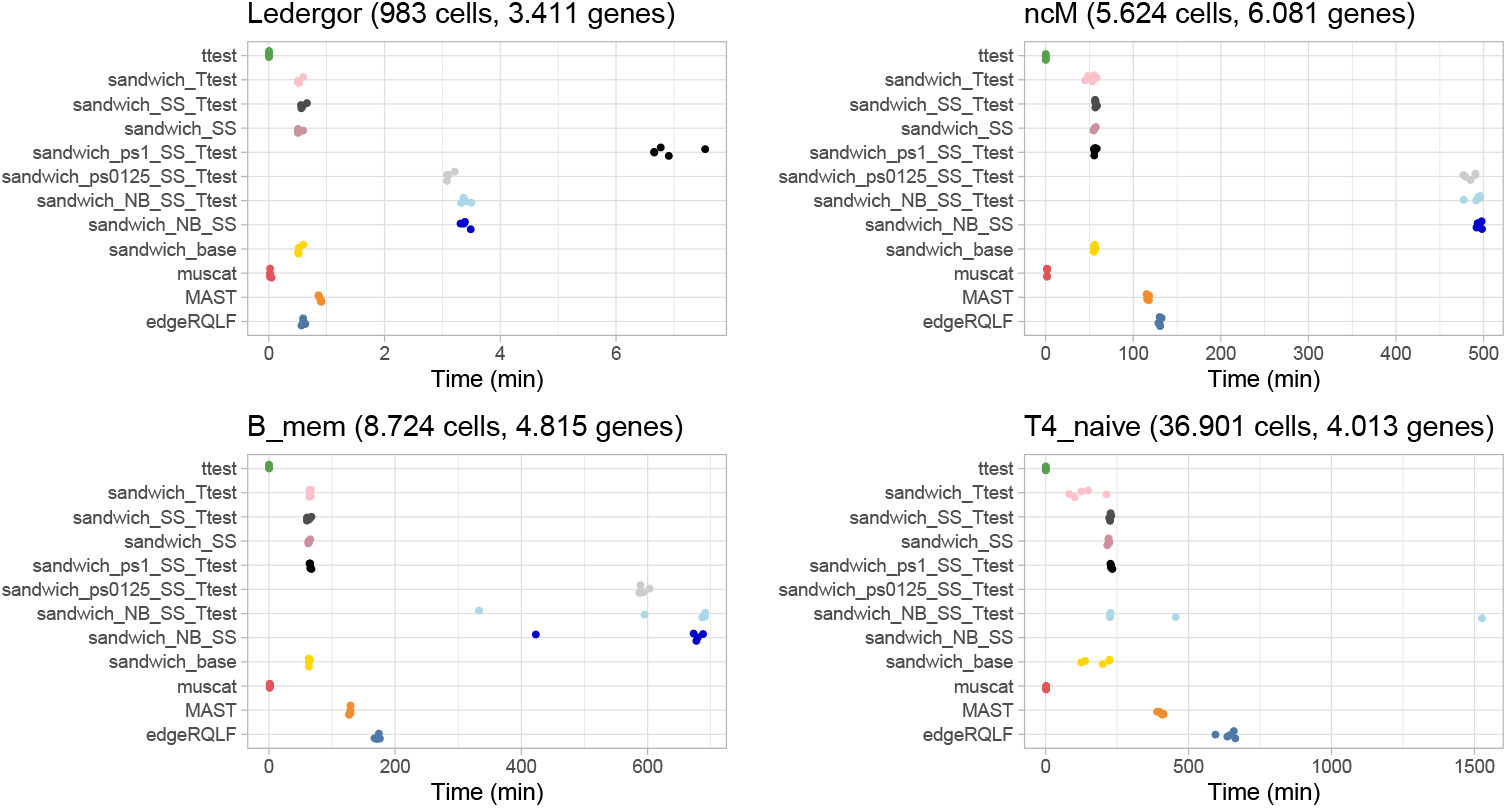
Running time of all DE methods on the different mock simulation datasets. Note that each method will have five running times for each dataset, i.e., one run time for each data replicate. **Panel A:** Running time for the mock simulations based on the Ledergor data. **Panel B:** Running time for the mock simulations based on the memory B-cells of the Lupus data. **Panel C:** Running time for the mock simulations based on the non-classical myeloid cells of the Lupus data. **Panel D:** Running time for the mock simulations based on the T4naive cells of the Lupus data.

Across all datasets, the t-test procedure and *muscat* had the fastest running times. This is unsurprising, as the t-test procedure is a very simple method, and as *muscat* performs a DE test on pseudobulk aggregated data, thereby greatly reducing data dimensionality (10 observations for the Ledergor dataset, 44 observations for the Lupus dataset). Next, all GEE methods that assume a Poisson distribution for the data and that do not incorporate a pseudo-count have a good scalability, running for approximately 1 hour on the Lupus Bmem and ncM datasets and below 4 hours on the large Lupus T4 naive dataset with 36.901 observations. MAST and edgeRQLF were slightly slower than these GEE methods, but still in the same ballpark. GEE methods that assume a Negative Binomial variance function (sandwich_NB_SS and sandwich_NB_SS_Ttest) display a considerably worse runtime than Poisson GEE methods. This is unsurprising, as the Negative Binomial variance function estimates an additional dispersion parameter to model a mean-variance relationship in the data. Due to their poor scalability, sandwich_NB_SS was not included for the Lupus T4 naive dataset. Finally, the behavior of the run times of Poisson GEE methods that incorporate a pseudo-count (sandwich_ps1_SS_Ttest and sandwich_ps0125_SS_Ttest) is inconsistent across datasets. sandwich_ps1_SS_Ttest runs approximately 8 times slower than sandwich_SS_Ttest (i.e., the same method without pseudo-count) on the Ledergor dataset. However, for the Lupus datasets, this decrease in scalability w.r.t. sandwich_SS_Ttest was not observed. sandwich_ps0125_SS Ttest runs approximately 4 times slower than sandwich_SS_Ttest on the Ledergor dataset, and 8 times slower on the Lupus Bmem and ncM datasets. Given its poor scalability on Lupus Bmem and ncM datasets, sandwich_ps0125_SS_Ttest was excluded for the analysis on the Lupus T4 naive dataset.

Note that for this scalability evaluation, all methods were run on a single core, i.e., without parallel execution. As such, the runtime of all methods except for edgeRQLF, *muscat* and MAST could be readily reduced by running fitting the models in parallel across multiple cores.

### Performance evaluation

We assess the performance of the five main DE methods on simulated datasets with ground truth DE signal. As in the previous paragraph, the simulated datasets are generated based on the Ledergor dataset [23] and the Lupus dataset [25]. However, here the simulation framework introduces DE signal for 5% of the genes, thus allowing for evaluating the DE methods both in terms of type I error control and statistical power, i.e., their specificity and sensitivity. We refer to Section Simulations with DE signal for more details on the setup of these simulation studies.

Fig 4 displays the benchmark results on simulated data based on the Ledergor dataset (panel A) and three cell types from the Lupus dataset, i.e., memory B cells (B mem, panel B), non-classical myeloid cells (ncM, panel C) and T4 naive cells (T4 naive, panel D). The Ledergor dataset consists of 5 samples per treatment group, whereas the Lupus dataset consists of 22 samples in each treatment group for each cell type. The analysis was performed five times per dataset, each time with a different randomly selected subset of samples per treatment group, and results were averaged across replicate analyses. Each curve visualizes the performance of a DE method by displaying its sensitivity (true positive rate, TPR) versus its specificity (false discovery rate, FDR). We refer to Section Performance measures for a detailed description of how these measures were computed.

**Fig 4.**
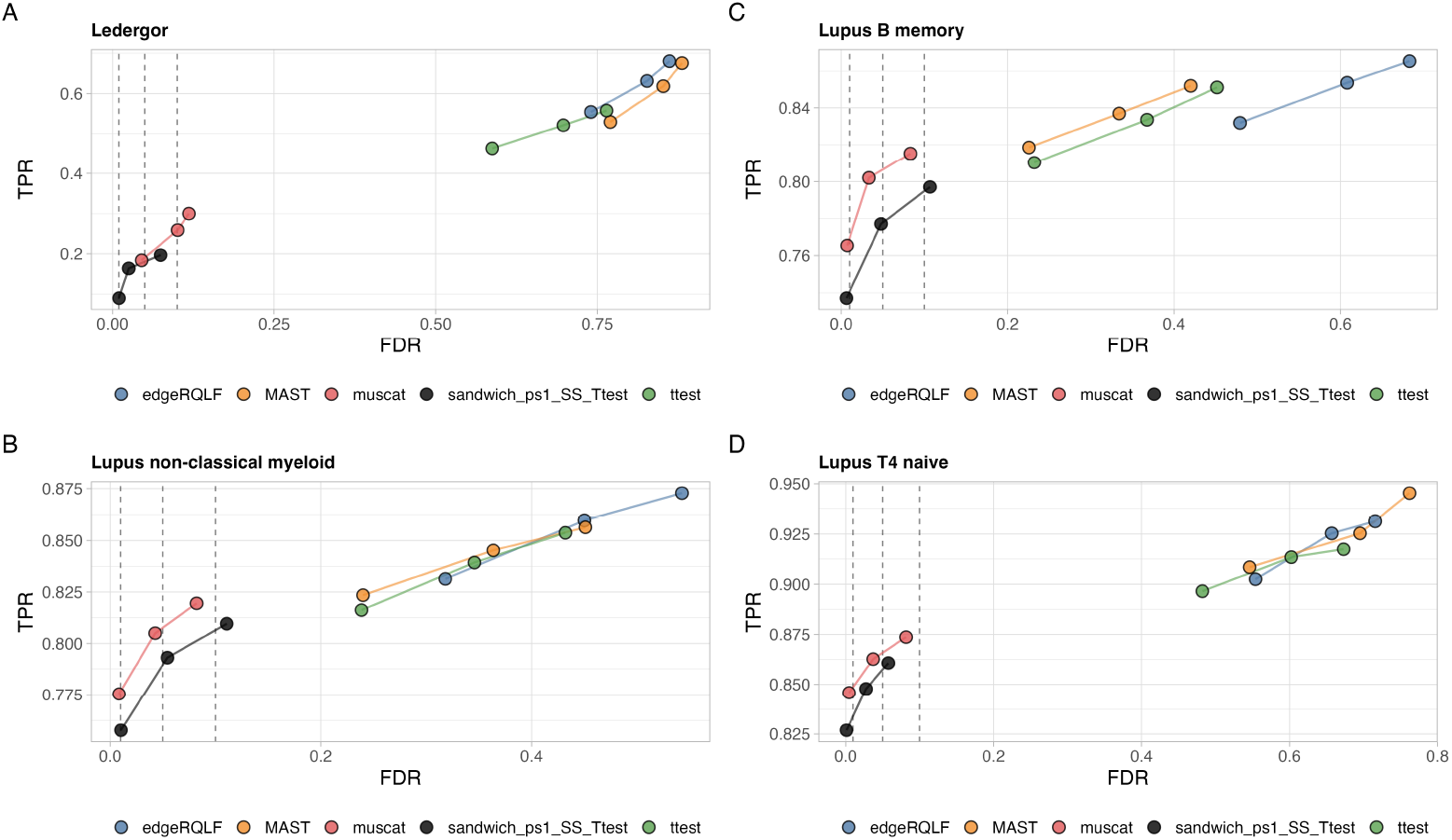
Performance benchmark of the different DE methods on simulated data using FDR-TPR curves. Each curve visualizes the performance of each method by displaying the sensitivity of the method (true positive rate, TPR) with respect to the false discovery rate (FDR). The curves display averages over 5 replicates for each simulated dataset. The three circles on each curve represent working points when the FDR level is set at nominal levels of 1%, 5% and 10%, respectively. **Panel A:** Performance evaluation on the Ledergor data (5 vs. 5 comparison). **Panel B:** Performance evaluation on the Lupus data, non-classical myeloid cell type (22 vs. 22 comparison). **Panel C:** Performance evaluation on the Lupus data, B memory cell type (22 vs. 22 comparison). **Panel D:** Performance evaluation on the Lupus data, T4 naive cell type (22 vs. 22 comparison).

For all datasets, *muscat* adequately controls the type I error and has a high sensitivity. While the sandwich method also provides good type I error control, its sensitivity is considerably lower than that of *muscat*. The three canonical single-cell DE methods (edgeRQLF, MAST and ttest) have poor specificity and, when evaluated at the same nominal FDR as *muscat*, a lower sensitivity than *muscat*. Note that the sensitivity for all methods is highest for the T4 naive cell type. However, the type I error control of the canonical single-cell DE methods is worst for this cell type. This is likely because this is the cell type with the highest number of cells per patient, i.e., the cell type with the largest level of pseudoreplication. Finally, the sensitivity is much higher for the three cell types of the Lupus data compared to the Ledergor data (note the difference in the y-axis range). As discussed in the next paragraph, this can in part be explained by the larger number of patients in the Lupus data (22 vs. 22 comparison) than in the Ledergor data (5 vs. 5 comparison).

Supplementary Fig S5 provides an alternative representation of the specificity and sensitivity of the different DE methods for the simulated data based on three cell types from the Lupus dataset. This visualization allows for assessing the FDP (y-axis) and TPR (dot sizes) for each cell type (dot shape), with each of the five replicates per cell type. This plot clearly shows that the canonical single-cell DE methods report too many false positives across all replicates for all cell types, and for the T4 naive cell type in particular (diamond dot shape).

Performance evaluation curves for all DE methods discussed in Sections DE methods from the existing literature and Generalized Estimation Equations for multi-patient scRNA-seq DE testing on the different simulated datasets are displayed in Supplementary Fig S6-S9. Across all four simulation datasets, we observe that all GEE-based workflows display similar relationships between FDR and TPR, but that only those methods that (1) use a small sample size correction and (2) rely on a t-test for the inference are able to accurately control the FDR at the desired level, which is in line with our mock analysis results (see Section Mock analysis). Furthermore, we observe that the performance of the methods with a pseudo-count (sandwich_ps1_SS_Ttest and sandwich_ps0125_SS_Ttest) are always higher or on par with that of the sandwich_SS_Ttest method that does not use a pseudo-count. Note that the sandwich_ps0125_SS_Ttest method was not included in the Lupus dataset simulation due to its poor scalability (as displayed in Fig 3).

### Influence of the number of subjects

To investigate the influence of the number of subjects on the sensitivity and specificity of the different DE methods, the Lupus data (ncM cell type) was downsampled to create comparisons with sample sizes of 5 vs. 5, 10 vs. 10, 15 vs. 15 and 22 vs. 22 samples. The downsampling was performed five times per sample size, each time with a different randomly selected subset of samples. The sensitivity and specificity of *muscat* and the sandwich method are displayed in Fig 5.

**Fig 5.**
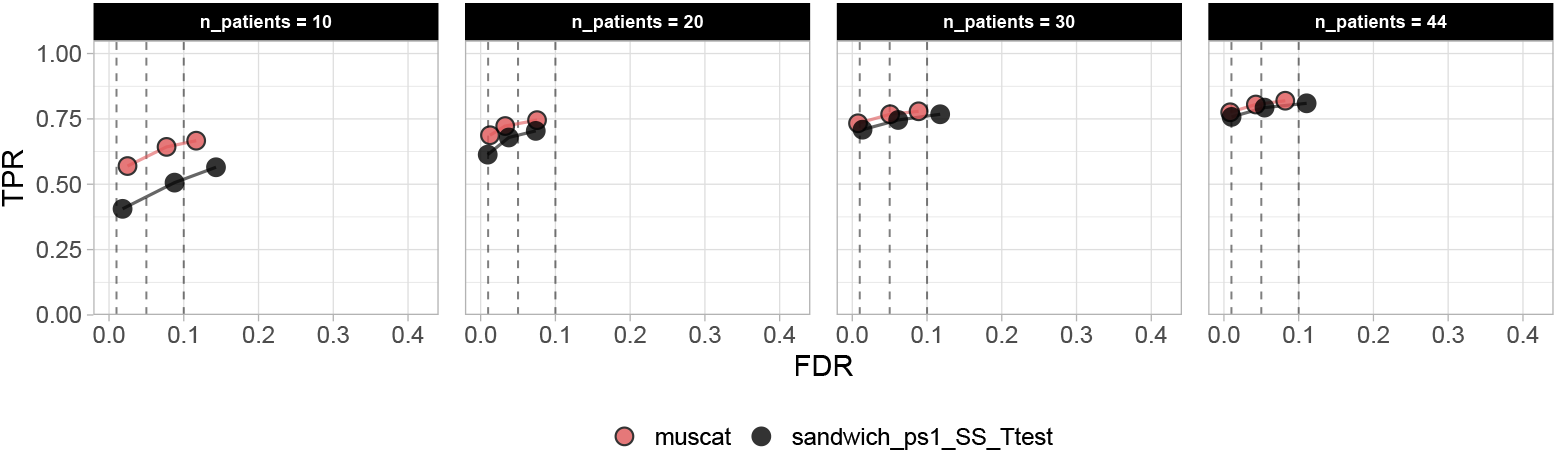
Influence of sample size on the performance of DE methods. The Lupus data (ncM cell type) was downsampled to create comparisons with sample sizes of 5 vs. 5, 10 vs. 10, 15 vs. 15 and 22 vs. 22 samples, as indicated in the header of the respective panels. Each curve visualizes the performance of a DE method by displaying sensitivity (true positive rate, TPR) with respect to the false discovery rate (FDR). Each curve is an average over 5 replicates for each simulated dataset. The three circles on each curve represent working points when the FDR level is set at nominal levels of 1%, 5% and 10%, respectively.

As expected, the sensitivity of both *muscat* and the sandwich method increases with increasing sample size (Fig 5). While the performance of the sandwich method is consistently lower than that of *muscat*, the difference is largest for the comparison with the smallest sample size (Fig 5, left panel). From the literature, it is known that GEE models have reduced power when the number of independent replicates is low [18]. Note that both methods properly control the type I error across the different sample sizes. While GEE models typically become overly liberal when the number of independent replicates is low [18, 20–22], this effect is not observed here, which is likely due to the small-sample size bias correction.

## Discussion

While multi-patient scRNA-seq datasets are becoming more and more common, there are still only a handful of DE methods that account for the within-subject cell-cell correlation inherent to such data. Hence, most current DE analysis workflows ignore pseudoreplication, which eventually results in overly liberal predictions [5]. In this manuscript, we have aimed to perform an unbiased comparison of various strategies to deal with pseudoreplication in DE analyses. Specifically, we benchmarked the performance of GEE models, a modeling strategy that has been mostly overlooked in previous studies. GEEs directly estimate the within-patient correlation structure of the data at the single-cell level, thereby omitting the need for aggregating the single-cell level data into pseudobulk profiles. As opposed to a previous study that benchmarked Gaussian GEEs on log-normalized scRNA-seq data [6], we used a Poisson GEE on the raw count data, thus respecting the count nature of the data.

Our benchmarks show that GEEs do not outperform *muscat* for analyses with simple two-group designs (Section Performance evaluation). The performance of GEEs is particularly lower for datasets with small sample sizes (Section Influence of the number of subjects), as the robustness of the sandwich estimator is an asymptotic property that requires a large number of subjects and a small number of repeated measurements (e.g. cells or cell types) per subject [15, 18]. While scRNA-seq datasets that profile a large number of subjects are becoming more and more common [1], these subjects are often distributed over a large number of groups or conditions. As such, the number of samples in each stratum often still is relatively small. For instance, while the Lupus dataset that we used in our benchmarks profiled 261 donors in total [25], the largest biologically homogeneous subgroup only contained 44 donors (see Section Mock simulation). In addition to having a higher statistical power, *muscat* is much faster to fit because the data are summarized at the pseudobulk level.

To avoid that some DE methods are favored over others because they use the same underlying model as scRNA-seq data simulation framework, our goal was to simulate data using a simulator that requires as little assumptions as possible. However, there is currently a lack of simulators that properly capture the characteristics of real scRNA-seq data [27]. This is particularly true for multi-subject scRNA-seq data, where the within-sample cell-cell correlations should be taken into account by the simulation framework. Therefore, we implemented a non-parametric simulator that does not rely on any model assumptions and retains as much of the original data as possible. The implementation of this minimal-assumptions simulator is available at https://github.com/milanmlft/swapper.

One relevant class of models that we did not consider in this manuscript are generalized linear mixed models (GLMMs), which can deal with pseudoreplication by including a random subject effect. We excluded GLMMs from our benchmark because previous benchmarks showed that GLMMs are computationally expensive and do not outperform pseudobulk methods [5, 7].

In conclusion, our benchmarks suggest that when the interest lies in testing for DE, i.e., changes in the average gene expression, pseudobulk-level analyses are preferred over single-cell level analyses. However, single-cell level methods are still useful if the goal is to assess differences in other moments of the expression distribution (not the mean), e.g., when studying higher order moments like variability, skewness, or curtosis. When modeling DE in multi-patient scRNA-seq datasets, our benchmarks clearly indicate that within-patient correlation should be acknowledged in order to avoid returning a massive number of false positive predictions.

## Acknowledgements

This work was supported by grants from Ghent University Special Research Fund (BOF20/GOA/023) (L.C.), Research Foundation Flanders (FWO G062219N) (J.G., L.C.) and (FWO SB fellowship No. 3S037119) (J.G.).

## Supporting Information

**S1 P-value density plots for all DE methods on null simulations based on the Ledergor dataset**. The different line types indicate the p-value densities for the five different data replicates.

**S2 P-value density plots for all DE methods on null simulations based on the B memory cell type of the Lupus dataset**. The different line types indicate the p-value densities for the five different data replicates.

**S3 P-value density plots for all DE methods on null simulations based on the T4 naive cell type of the Lupus dataset**. The different line types indicate the p-value densities for the five different data replicates.

**S4 P-value density plots for all DE methods on null simulations based on the B memory cell type of the Lupus dataset**. The different line types indicate the p-value densities for the five different data replicates.

**S5 Performance benchmark of the different DE methods on simulated data using scatterplots**. Each point represents a the observed FDP of a specific simulation replicate at a nominal FDP level of 0.05. The points are sized according to the TPR. **Panel A:** Performance evaluation on three different cell types of the Ledergor data. **Panel B:** Performance evaluation on three different cell types of the Lupus data.

**S6 Performance benchmark of all the different DE methods on the simulated data based on the Ledergor dataset**. Each curve visualizes the performance of each method by displaying the sensitivity of the method (true positive rate, TPR) with respect to the false discovery rate (FDR). Each curve displays the average profile over 5 replicates for each simulated dataset. The three circles on each curve represent working points when the FDR level is set at nominal levels of 1%, 5% and 10%, respectively.

**S7 Performance benchmark of all the different DE methods on the simulated data based on the Lupus ncM cell type dataset**. Each curve visualizes the performance of each method by displaying the sensitivity of the method (true positive rate, TPR) with respect to the false discovery rate (FDR). Each curve displays the average profile over 5 replicates for each simulated dataset. The three circles on each curve represent working points when the FDR level is set at nominal levels of 1%, 5% and 10%, respectively.

**S8 Performance benchmark of all the different DE methods on the simulated data based on the Lupus B memory cell type dataset**. Each curve visualizes the performance of each method by displaying the sensitivity of the method (true positive rate, TPR) with respect to the false discovery rate (FDR). Each curve displays the average profile over 5 replicates for each simulated dataset. The three circles on each curve represent working points when the FDR level is set at nominal levels of 1%, 5% and 10%, respectively.

**S9 Performance benchmark of all the different DE methods on the simulated data based on the Lupus T4 naive cell type dataset**. Each curve visualizes the performance of each method by displaying the sensitivity of the method (true positive rate, TPR) with respect to the false discovery rate (FDR). Each curve displays the average profile over 5 replicates for each simulated dataset. The three circles on each curve represent working points when the FDR level is set at nominal levels of 1%, 5% and 10%, respectively.

