## Supplemental Text and Figures for "Strategies for addressing pseudoreplication in multi-patient scRNA-seq data"

### Supporting Information

#### Supplementary figures

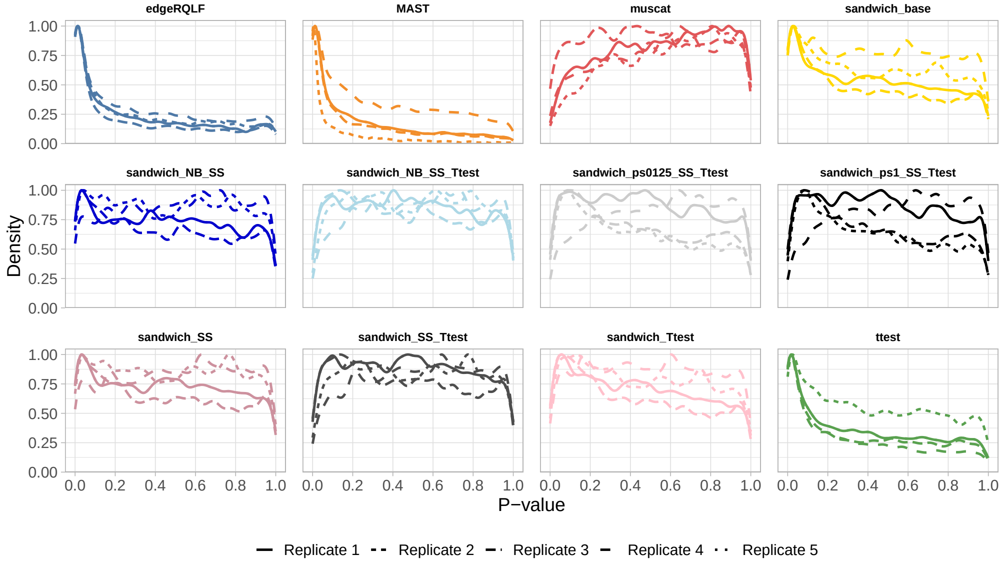


**Figure S1. P-value density plots for all DE methods on null simulations based on the Ledergor dataset.** The different line types indicate the p-value densities for the five different data replicates.

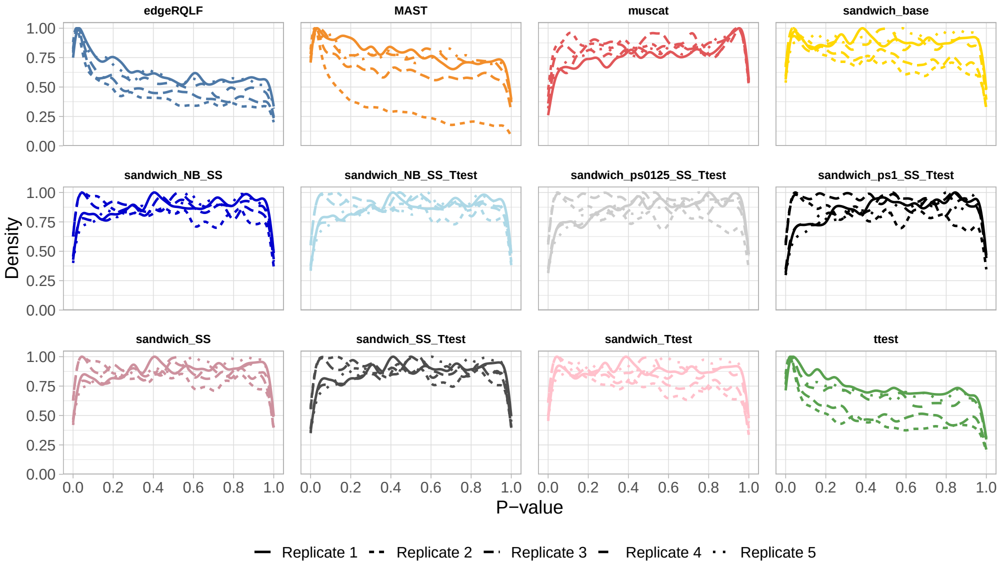


**Figure S2. P-value density plots for all DE methods on null simulations based on the B memory cell type of the Lupus dataset.** The different line types indicate the p-value densities for the five different data replicates.

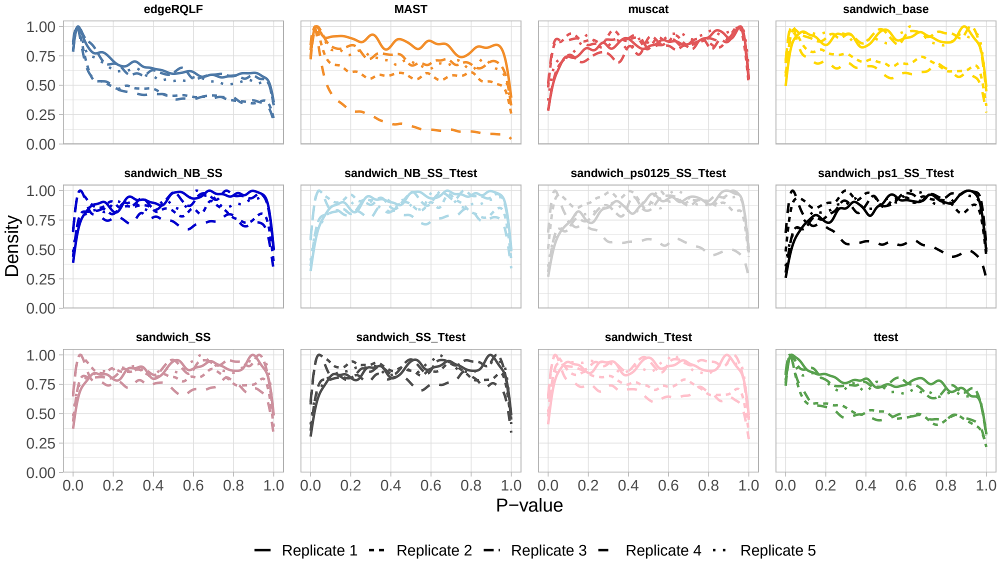


**Figure S3. P-value density plots for all DE methods on null simulations based on the non-classical myeloid (ncM) cell type of the Lupus dataset.** The different line types indicate the p-value densities for the five different data replicates.

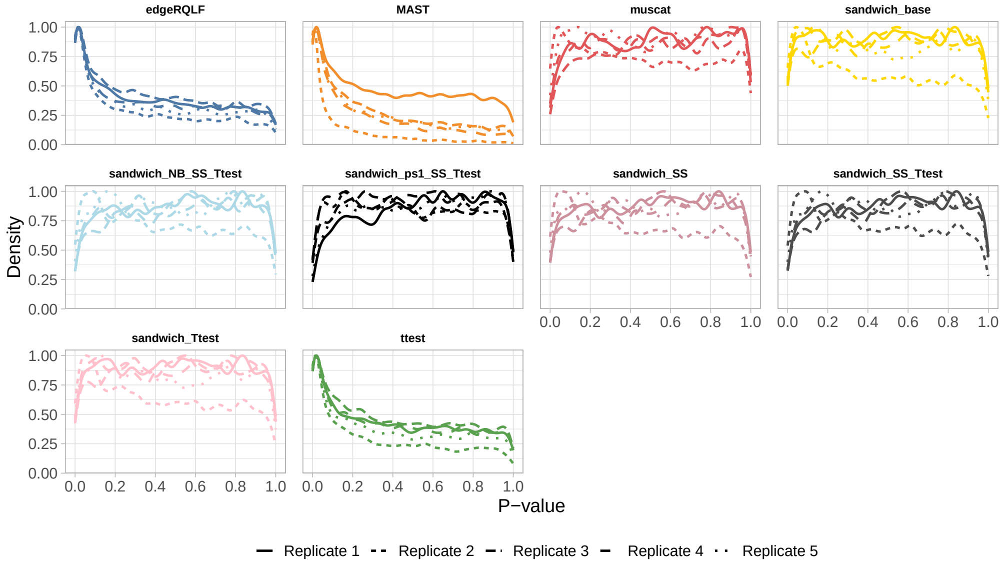


**Figure S4. P-value density plots for all DE methods on null simulations based on the T4 naive cell type of the Lupus dataset.** The different line types indicate the p-value densities for the five different data replicates.

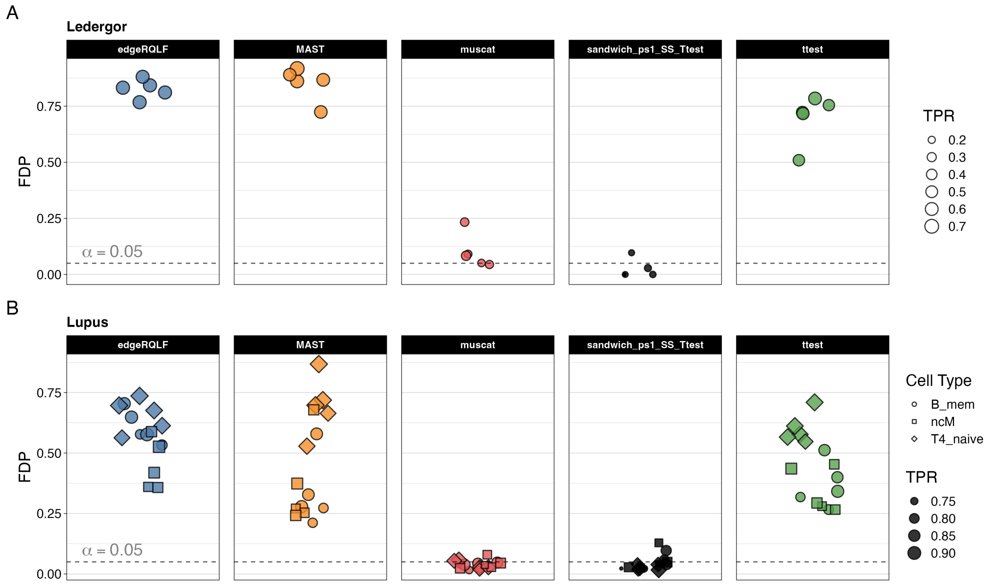


**Figure S5. Performance benchmark of the different DE methods on simulated data using scatterplots.** Each point represents a the observed FDP of a specific simulation replicate at a nominal FDP level of 0.05. The points are sized according to the TPR. **Panel A:** Performance evaluation on three different cell types of the Ledergor data. **Panel B:** Performance evaluation on three different cell types of the Lupus data.

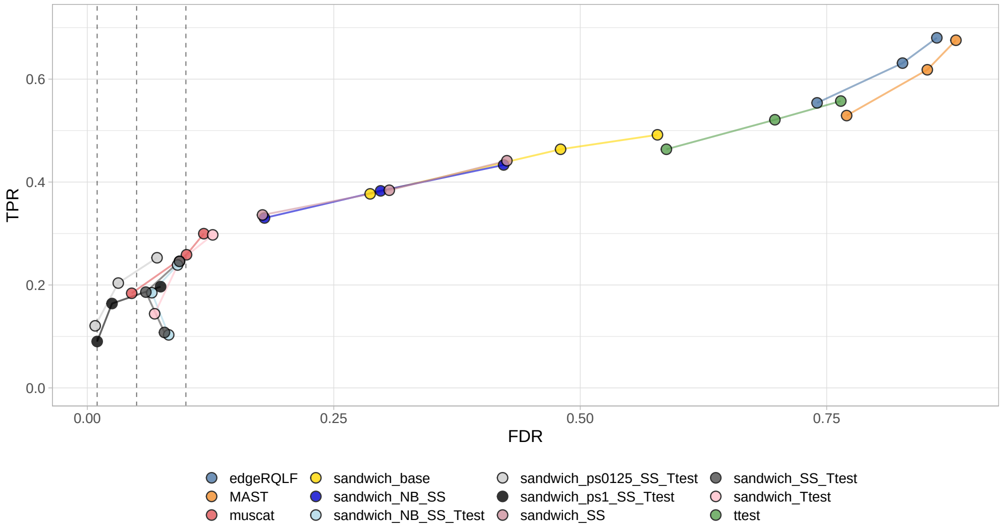


**Figure S6. Performance benchmark of all the different DE methods on the simulated data based on the Ledergor dataset.** Each curve visualizes the performance of each method by displaying the sensitivity of the method (true positive rate, TPR) with respect to the false discovery rate (FDR). Each curve displays the average profile over 5 replicates for each simulated dataset. The three circles on each curve represent working points when the FDR level is set at nominal levels of 1%, 5% and 10%, respectively.

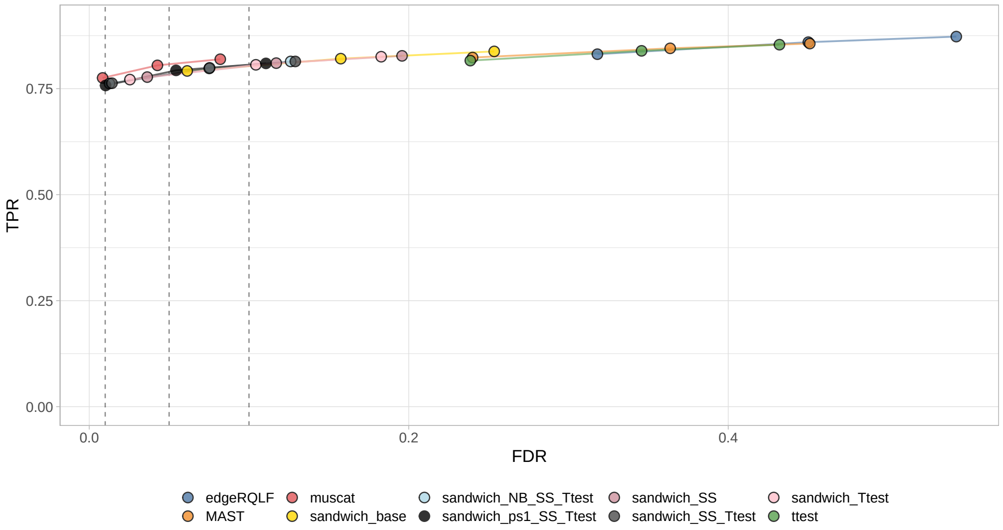


**Figure S7. Performance benchmark of all the different DE methods on the simulated data based on the Lupus ncM cell type dataset.** Each curve visualizes the performance of each method by displaying the sensitivity of the method (true positive rate, TPR) with respect to the false discovery rate (FDR). Each curve displays the average profile over 5 replicates for each simulated dataset. The three circles on each curve represent working points when the FDR level is set at nominal levels of 1%, 5% and 10%, respectively.

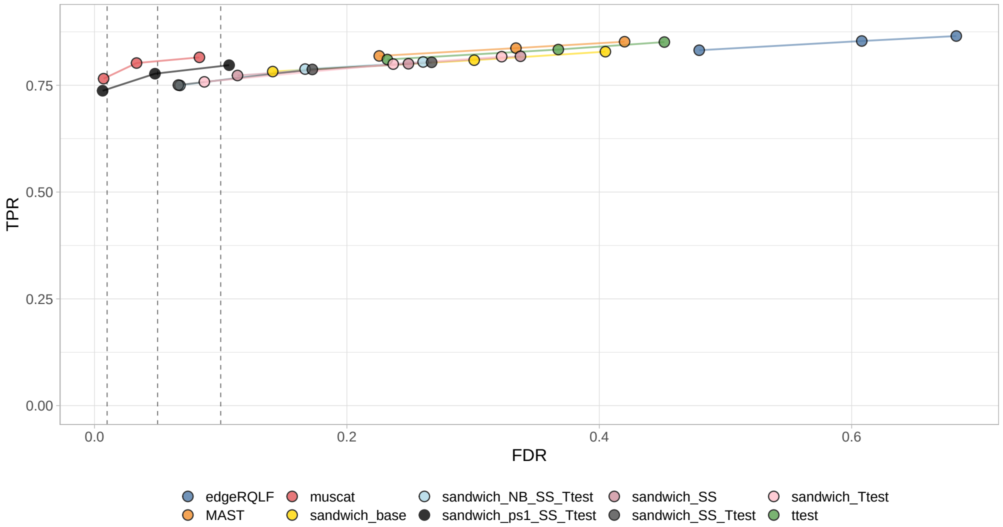


**Figure S8. Performance benchmark of all the different DE methods on the simulated data based on the Lupus B memory cell type dataset.** Each curve visualizes the performance of each method by displaying the sensitivity of the method (true positive rate, TPR) with respect to the false discovery rate (FDR). Each curve displays the average profile over 5 replicates for each simulated dataset. The three circles on each curve represent working points when the FDR level is set at nominal levels of 1%, 5% and 10%, respectively.

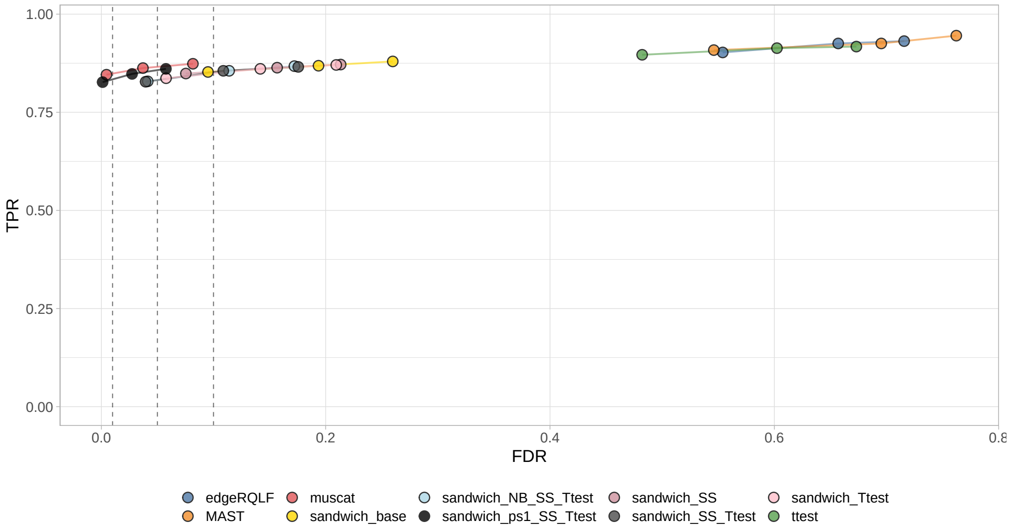


**Figure S9. Performance benchmark of all the different DE methods on the simulated data based on the Lupus T4 naive cell type dataset.** Each curve visualizes the performance of each method by displaying the sensitivity of the method (true positive rate, TPR) with respect to the false discovery rate (FDR). Each curve displays the average profile over 5 replicates for each simulated dataset. The three circles on each curve represent working points when the FDR level is set at nominal levels of 1%, 5% and 10%, respectively.

#### Technical note

Since the advent of the micro-array and bulk RNA-seq technologies, Generalized Linear Models (GLMs) have been the flagship framework for modeling RNA abundance data. Some popular GLM-based DE methods directly model the discrete RNA abundance measurements using count distributions like the Negative Binomial distribution (e.g., DESeq2 [1] and edgeR [2]), whereas other methods first normalize the RNA abundance counts to allow for Gaussian linear modeling (e.g. limma [3], limma-voom [4] and MAST [5]). One disadvantage of GLMs is that they cannot readily accommodate the hierarchical correlation structure commonly observed in multi-sample scRNA-seq data. Indeed, when multiple cells from the same subject are profiled, the cell-level expression profiles can no longer be considered independent. This within-subject cell-cell correlation, also referred to as pseudoreplication, must be accounted for in the downstream statistical modeling [6–8]. One way of overcoming pseudoreplication is to aggregate the data at the pseudobulk level [7]. Another possibility is to explicitly model the within-subject correlation at the single-cell level. This can either be achieved using Generalized Linear Mixed Models (GLMMs), also referred to as Mixed-effect models, or by Generalized Estimation Equestions (GEEs).

Several implementations of GLMMs for the analysis of scRNA-seq data exist [9–11]. However, canonical GLMMs scale poorly to datasets with a large number of observations [7, 8]. This issue has been reduced to some extent by the highly efficient GLMM implementations by Brooks *et al.* (2017) [9] and He *et al.* (2021) [11]. The performance of canonical GLMMs has already been extensively benchmarked by Squair *et al.* (2021) [8] and Murphy and Skene (2022) [12]. These benchmarks both indicated that the performance of GLMMs was consistently lower than the performance of pseudobulk methods on datasets with a large number of cells, and on par with pseudobulk methods for small datasest (25-50 cells). As such, we do not consider GLMMs in this benchmark.

GEEs provide an alternative strategy for modeling data with repeated measurements. They were first developed by Liang and Zeger (1986) [13] in the context of longitudinal data analysis (repeated measurements within a subject along time) for a binary outcome. However, GEEs are not limited to these settings. They provide a general statistical framework for extending GLMs to repeated measurements data. The model parameters from GEEs have a marginal or population-average interpretation (below we will show how this follows from the GEE model specification).

We here formulate the specification of the GEE model. As an example, we take an scRNA-seq dataset with multiple subjects and multiple cells per subjects. Without loss of generality, we will consider a single gene $g$, and henceforth drop the subscript $g$. Let:

$$\mathbf{Y_i} = (Y_{i1}, ..., Y_{ij}, ..., Y_{in_i}) \tag{1}$$

be a $n_i \times 1$ column vector of responses (measured gene abundances) for cells $j = 1, ..., n_i$ of subject $i = 1, ..., N$ and

$$\mathbf{X_{ij}} = (X_{ij1}, X_{ij2}, ..., X_{ijp}) \tag{2}$$

a $p \times 1$ column vector of between-subject and within-subject covariates potentially associated with the response $Y_{ij}$.

GEEs can then be described using a "three-way" specification, specifying (i) the expectation of the response conditional on the fixed effects $\boldsymbol{X_{ij}^T}$, (ii) the conditional variance of $Y_{ij}$ given $\boldsymbol{X_{ij}^T}$ and (iii) the conditional within-subject association among the vector of repeated responses [13, 14]:

| Correlation structure | $\mathrm{Corr}(Y_{ij}, Y_{ik})$ | Sample matrix |
|---|---|---|
| Independent | $\mathrm{Corr}(Y_{ij}, Y_{ik}) = \begin{cases} 1 & j = k \\ 0 & j \neq k \end{cases}$ | $\begin{pmatrix} 1 & 0 & 0 \\ 0 & 1 & 0 \\ 0 & 0 & 1 \end{pmatrix}$ |
| Exchangeable | $\mathrm{Corr}(Y_{ij}, Y_{ik}) = \begin{cases} 1 & j = k \\ \alpha & j \neq k \end{cases}$ | $\begin{pmatrix} 1 & \alpha & \alpha \\ \alpha & 1 & \alpha \\ \alpha & \alpha & 1 \end{pmatrix}$ |
| Unstructured | $\mathrm{Corr}(Y_{ij}, Y_{ik}) = \begin{cases} 1 & j = k \\ \alpha_{jk} & j \neq k \end{cases}$ | $\begin{pmatrix} 1 & \alpha_{12} & \alpha_{13} \\ \alpha_{21} & 1 & \alpha_{23} \\ \alpha_{31} & \alpha_{32} & 1 \end{pmatrix}$ |

**Figure S10. Summary of commonly used "working" correlation structures for GEE.** Note that many other correlation structures exist. We only show the three structures that are most relevant in the setting of non-longitudinal data, e.g., scRNA-seq data that are not acquired from time-course experiments. Figure adapted from [15].

$$\begin{aligned} (i) &\quad E(Y_{ij}|\mathbf{X_{ij}}) = g(\mu_{ij}) = \eta_{ij} = \boldsymbol{X_{ij}^T\beta} \\ (ii) &\quad Var(Y_{ij}|\mathbf{X_{ij}}) = \phi\nu(\mu_{ij}) \\ (iii) &\quad \mathbf{V_i} = \mathbf{A_i^{1/2}}Corr(\mathbf{Y_i})\mathbf{A_i^{1/2}} = \mathbf{A_i^{1/2}}\boldsymbol{R_i(\alpha)}\mathbf{A_i^{1/2}} \end{aligned} \tag{3}$$

Component (i) relates the expectation of the response with a linear predictor of covariates through a known link function $g(.)$. Note that this specification coincides with that of the mean component of a canonical GLM. In both cases, population-averaged or marginal effects are obtained. Indeed, the estimator of the mean model parameters of the GEE is conditional only on the fixed effects $\boldsymbol{X_{ij}^T}$, and not conditional on any random effects.

Component (ii) describes the mean-variance relationship, i.e., the relationship between the variance of $Y_{ij}$ given the covariates $\mathbf{X_{ij}}$ according to a known variance function $\nu_i(.)$, and an additional scale or dispersion parameter $\phi$. Note that in the most general formulation, which is displayed here, a single dispersion parameter $\phi$ is estimated. In principle, $\phi$ can be estimated separately for different covariate levels, e.g., $\phi$ can be cell type specific.

Note that component (i) and component (ii) specify only the first two moments of the distribution of $Y_{ij}$. Hence, GEEs are semi-parametric models.

Component (iii) explicitly describes the within-subject correlation conditional on the covariates $\mathbf{X_{ij}}$, as a function of the conditional means $\mu_{ij}$ and an additional set of within-subject association parameters $\alpha$. More specifically, $\mathbf{A_i}$ is a diagonal matrix with $Var(Y_{ij}|\mathbf{X_{ij}}) = \phi\nu(\mu_{ij})$ along the diagonal. As such, $\mathbf{A_i^{1/2}}$ is a diagonal matrix of the standard deviations. $Corr(\mathbf{Y_i}) = \boldsymbol{R(\alpha)}$ is a matrix describing the correlation among repeated measurements within subject $i$ as a function of parameters $\alpha$. Fig S10 displays three parametrizations of the within-subject correlation. Under independence, the repeated measurements within subject $i$ are considered independent from one another, i.e., $\alpha = 0$. The exchangeable correlation structure, sometimes also referred to as compound symmetry, assumes a constant pairwise correlation between all cells of subject $i$. Lastly, the unstructured correlation assumes a different value for $\alpha$ between each pair of cells from subject $i$. Finally, $\mathbf{V_i}$ is typically referred to as the "working" covariance matrix, which is allowed to be misspecified, as shown below (Eq 7).

The GEE estimator of the marginal mean model parameters are obtained by minimizing the objective function

$$\sum_{i=1}^{N} [\boldsymbol{Y_i} - \boldsymbol{\mu_i}]^{\boldsymbol{T}} \; \boldsymbol{V_i^{-1}} \; [\boldsymbol{Y_i} - \boldsymbol{\mu_i}] \tag{4}$$

with $\boldsymbol{\mu_i} = \boldsymbol{\mu_i(\beta)} = g^{-1}(\mathbf{X_{ij}}\boldsymbol{\beta})$ a column vector of the mean responses. It can be shown that minimizing Eq 4 requires solving the set of generalized estimation equations [13]

$$\sum_{i=1}^{N} \mathbf{D_i^T} \mathbf{V_i^{-1}} (\mathbf{Y_i} - \boldsymbol{\mu}_\mathbf{i}) = 0 \tag{5}$$

with $\mathbf{V_i^{-1}}$ the "working" covariance matrix specified in component (iii) of Eq 3, and $\mathbf{D_i} = \frac{\delta \boldsymbol{\mu_i}}{\delta \boldsymbol{\beta}}$ the matrix of derivatives of $\boldsymbol{\mu_i}$ with respect to $\boldsymbol{\beta}$.

Because the GEE (Eq 5) depends on both $\beta$ and $\alpha$, an iterative two-stage estimation procedure is required. In the first stage, based on the current estimates for $\alpha$, $\phi$ and $V_i$, an updated estimate of $\beta$ is obtained by solving Eq 5. In the second stage, updated estimates for $\alpha$ and $\phi$ are obtained based on standardized residuals $e_{ij} = (Y_{ij} - \widehat{\mu_{ij}})/\sqrt{v(\widehat{\mu_{ij}})}$, which are required to update $\mathbf{V_i}$.

The resulting solution $\hat{\boldsymbol{\beta}}$ for Eq 5 has some very appealing properties. First, $\hat{\boldsymbol{\beta}}$ is a consistent estimator for $\boldsymbol{\beta}$, with consistency depending on the correct specification of $\boldsymbol{\mu_i}$ but not of $\mathbf{V_i}$. As such, $\hat{\boldsymbol{\beta}}$ is consistent even if $\mathbf{V_i}$ is misspecified.

Secondly, in large samples, $\hat{\boldsymbol{\beta}}$ is multivariate normally distributed, i.e.,

$$\hat{\boldsymbol{\beta}} \sim \; MVN(\boldsymbol{\beta}, \mathbf{V}_{\boldsymbol{\beta}}^{\mathbf{R}}) \tag{6}$$

with mean $E(\hat{\boldsymbol{\beta}}) = \boldsymbol{\beta}$ and $\mathbf{V}_{\boldsymbol{\beta}}^{\mathbf{R}} = Cov(\hat{\boldsymbol{\beta}}) = BMB$, with

$$\begin{aligned} B &= (\sum_{i=1}^{N} \mathbf{D_i^T} \mathbf{V_i^{-1}} \mathbf{D_i})^{-1} \\ M &= \sum_{i=1}^{N} \mathbf{D_i^T} \mathbf{V_i^{-1}} Cov(\mathbf{Y_i}) \mathbf{V_i^{-1}} \mathbf{D_i} \end{aligned} \tag{7}$$

$\mathbf{V}_{\boldsymbol{\beta}}^{\mathbf{R}}$ can be estimated by plugging in the empirical values $\hat{D}_i$ and $\hat{V}_i$, and by replacing $Cov(\mathbf{Y_i})$ by $E[(\mathbf{Y_i} - \mathbf{X_i}\hat{\beta})(\mathbf{Y_i} - \mathbf{X_i}\hat{\beta})^T]$ [14]. $\widehat{\mathbf{V}_{\boldsymbol{\beta}}^{\mathbf{R}}}$ is commonly referred to as the sandwich variance estimator (also known as the empirical or robust variance estimator). The sandwich estimator is a robust estimator of the variance of $\hat{\boldsymbol{\beta}}$ in the sense that $M$ corrects for misspecification of the working covariance $\mathbf{V_i}$. However, note that the robustness property of the sandwich estimator only holds asymptotically, i.e., when (i) the number of subjects $N$ is relatively large, (ii) the number of repeated measurements per patient $n_i$ is relatively small and (iii) the number of repeated measurements is balanced across subjects. Note that for scRNA-seq data the reverse scenario, i.e., a low number of subjects, a high number of cells per subject and strong variability in the number of cells per subject, is often observed.

Note that a conventional GLM with a sandwich estimator is a special case of a GEE. Indeed, for GEEs that use an independent correlation structure (Fig S10), the estimation of the mean model parameters reduces to a Newton-Raphson algorithm that is analogous to that of a traditional GLM. Subsequently obtaining the variance with sandwich estimator makes inference robust towards the misspecification of the working correlation. In the remainder of this manuscript, we henceforth refer to GEEs with an independent correlation structure as "sandwich".
